# Precise and programmable C:G to G:C base editing in genomic DNA

**DOI:** 10.1101/2020.07.21.213827

**Authors:** Liwei Chen, Jung Eun Park, Peter Paa, Priscilla D. Rajakumar, Yi Ting Chew, Swathi N. Manivannan, Wei Leong Chew

## Abstract

Many genetic diseases are caused by single-nucleotide polymorphisms (SNPs). Base editors can correct SNPs at single-nucleotide resolution, but until recently, only allowed for C:G to T:A and A:T to G:C transition edits, addressing four out of twelve possible DNA base substitutions. Here we developed a novel class of C:G to G:C Base Editors (CGBEs) to create single-base genomic transversions in human cells. Our CGBEs consist of a nickase CRISPR-Cas9 (nCas9) fused to a cytosine deaminase and base excision repair (BER) proteins. Characterization of >30 CGBE candidates and 27 guide RNAs (gRNAs) revealed that CGBEs predominantly perform C:G to G:C editing (up to 90% purity), with rAPOBEC-nCas9-rXRCC1 being the most efficient (mean C:G to G:C edits at 15% and up to 37%). CGBEs target cytosine in WCW, ACC or GCT sequence contexts and within a precise two-nucleotide window of the target protospacer. We further targeted genes linked to dyslipidemia, hypertrophic cardiomyopathy, and deafness, showing the therapeutic potential of CGBE in interrogating and correcting human genetic diseases.

## Main text

Many human genetic diseases are caused by single-nucleotide polymorphism (SNPs), in which the disease and healthy alleles differ by a single DNA base. Base editors can correct these SNPs by converting the targeted DNA bases into another base in a controllable and efficient fashion. Current technology enables the conversion of C:G base pairs to T:A base pairs using cytosine base editors (CBEs) and A:T base pairs to G:C base pairs using adenine base editors (ABEs)^1–3^, which together represent half of all known disease-associated SNPs. CBEs and ABEs are also known to effect some C:G to G:C edits as byproducts^4,5^, but they cannot effect such transversions at efficiencies or purities necessary for therapeutic use, and hence the remaining half of these SNPs are not addressable by current base-editors^1,3^. Here, we developed a base editor capable of editing C:G to G:C (Fig. 1a), which opens up treatment avenues to half of all known SNP-associated human genetic diseases (Extended Data Table 1).

**Fig. 1.**
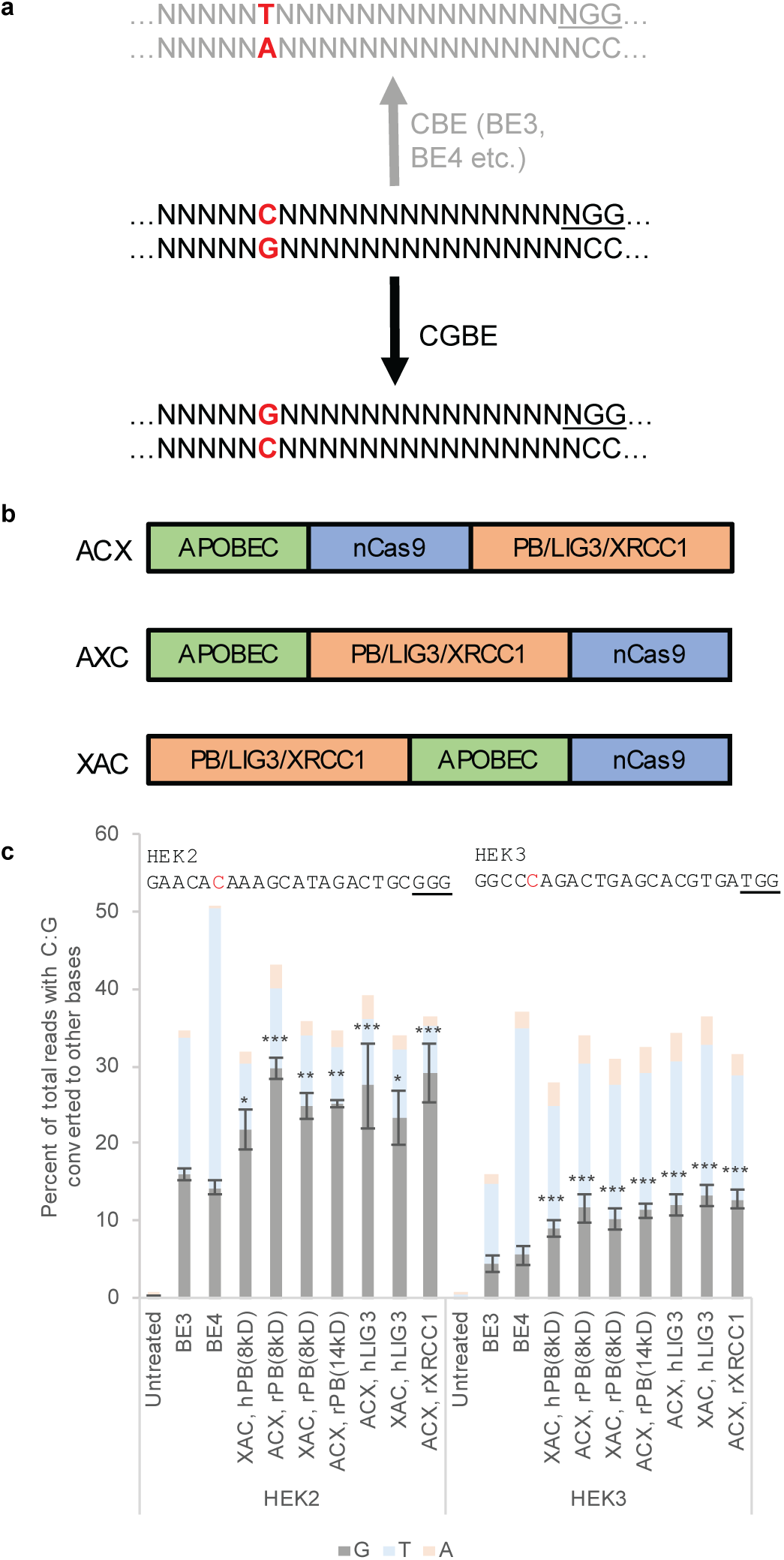
Initial screen of CGBE candidates for C:G to G:C editing. **(a)** CBEs like BE3 and BE4 predominantly convert C:G to T:A while CGBE aims to predominantly convert C:G to G:C. **(b)** CGBE candidates were designed in three orientations – ACX, AXC, and XAC, where X denotes the fused BER protein. **(c)** Seven candidates were selected for their high C:G to G:C editing at both HEK2 and HEK3. Targeted C’s are in red. PAMs are underlined. *p < 0.05; **p < 0.01; ***p < 0.001 (one-way ANOVA against ‘Untreated,’ Dunn-Šidák; n=3, mean ± standard error).

To effect C:G to G:C editing, we looked to leverage the cell’s innate base excision repair (BER) pathway, in which DNA polymerase β, DNA ligase III and XRCC1 are major players^6^. Previous work suggests that removal of uracil glycosylase inhibitor (UGI) from a CBE might lead to an increase in C:G to G:C levels^2^. UGI inhibits uracil DNA glycosylase (UDG), thereby inhibiting the downstream BER^1,2,7^. We speculate that the endogenous BER pathway – in the presence of a bound nCas9 – is linked to the observed C:G to G:C editing and that replacing UGI with BER proteins might increase such editing events. Using a version of CBE – BE3 – as a starting point, we removed UGI, instead fusing rAPOBEC-nCas9 with DNA ligase 3, DNA repair protein XRCC1, or the DNA binding and lyase domain of DNA polymerase β (PB)^8^.

Because the relative orientation of Cas9 fusions may affect the activity of the fusion, we built several versions of our CGBE candidates with the fused rAPOBEC and BER proteins at different orientations with respect to nCas9 (Fig. 1b). We then separately treated HEK293AAV cells with each candidate along with gRNAs designed to target genomic sites HEK2 and HEK3, both of which were previously used to characterize base editors^1^. As controls, we also treated cells with BE3 and BE4. Since no effective means of C:G to G:C base editing is currently known and BE3 has been observed to effect this reaction as a byproduct of C:G to T:A editing^4^, we used BE3 as a benchmark for our CGBEs.

High-throughput sequencing of the HEK2 site revealed that our CGBE candidates were able to edit C:G to both G:C and T:A. Out of the 31 candidates tested, 20 candidates showed an increased level of C:G to G:C editing relative to BE3 at position 6 within the protospacer (Extended Data Fig. 1a). The next closest available C was at position 4, where editing levels were lower (< 10%, Extended Data Fig. 1b), suggesting a narrower editing window than BE3^1^. In the best performing candidates, up to 29% C:G to G:C editing and 6% C:G to T:A editing were observed for the CGBE candidates, compared with 14% and 36% respectively for BE4, and 16% and 18% respectively for BE3.

Similarly, sequencing at the HEK3 site revealed editing of C:G to both G:C and T:A. 12 out of 31 candidates gave C:G to G:C editing levels at position 5 that were higher than that achieved by BE3, with the best performing candidates effecting up to 13% C:G to G:C editing compared with 4% for BE3 (Extended Data Fig. 2a). Despite an increase in C:G to T:A editing with CGBEs compared to BE3, a more than twofold increase in C:G to G:C editing with CGBEs led to an increase in the editing ratio of C:G to G:C compared to C:G to T:A. Compared to position 5, C:G to G:C editing at position 3 and position 4 are significantly lower (Extended Data Fig. 2b and 2c). From this pilot screen, we shortlisted seven of the best performing candidates for a secondary screen (Fig. 1c).

We further tested the CGBEs for C:G to G:C editing at four genomic sites known to be amenable to BE3-mediated editing – EMX1, HEK4, RNF2, and FANCF (Extended Data Fig. 3a). With CGBEs, C:G to G:C edits were efficiently induced (17-24%, compared to 8-10% with BE3) as the predominant product (up to 69% purity) at HEK4 and RNF2. At sites EMX1 and FANCF, up to 9% C:G to G:C editing was observed despite this not being the predominant edit, while BE3 effected up to 3% C:G to G:C editing. From this secondary screen, we chose ACX, rXRCC1 and ACX, rPB(8kD) for further characterization.

We next determined if target sequence context impacts editing efficiency. In characterizing the two CGBEs with eight other gRNAs – including dyslipidemia-associated gene ADRB2^9^, hearing loss-associated gene GJB2^10^, and hypertrophic cardiomyopathy-associated gene MYBPC3^11^ – we noticed that not only did CGBEs efficiently interrogate disease-associated genes, but they also gave higher levels of C:G to G:C editing at C’s immediately following an A/T (Extended Data Fig. 3b, Extended Data Fig. 4, and Extended Data Table 2). Hence, we suspected that the difference in editing efficiency between gRNAs might be due to the different motifs within which the targeted C is located (e.g. ACA at HEK2, CCA at HEK3; targeted C is underlined). To address such potential differences, we designed 16 guides targeting HEK2. These guides collectively contained all possible NCN motifs with targeted C’s at position 6 (Fig. 2a, Extended Data Fig. 3c). Sequencing revealed that the most readily edited motif was ACA, while the least edited motif was GCC. Up to 10% C:G to G:C editing was observed for ACA, ACC, ACT, GCT, TCA, and TCT. This observation applied to both ACX, rPB(8kD) and ACX, rXRCC1. From these results, we propose that the favored DNA motifs are WCW, ACC, and GCT (W is either A or T, Fig. 2b). This understanding is consistent with our observation that C:G to G:C editing is more efficient at HEK2, HEK4, and RNF2, which all had favored DNA motifs of ACA, ACT, and TCT, whereas EMX1, FANCF, and HEK3, with suboptimal DNA motifs of TCC and CCA, are less amenable to C:G to G:C editing. With a preference for WCW, ACC, and GCT, CGBEs most efficiently target 6 out of 16 possible motifs.

**Fig. 2.**
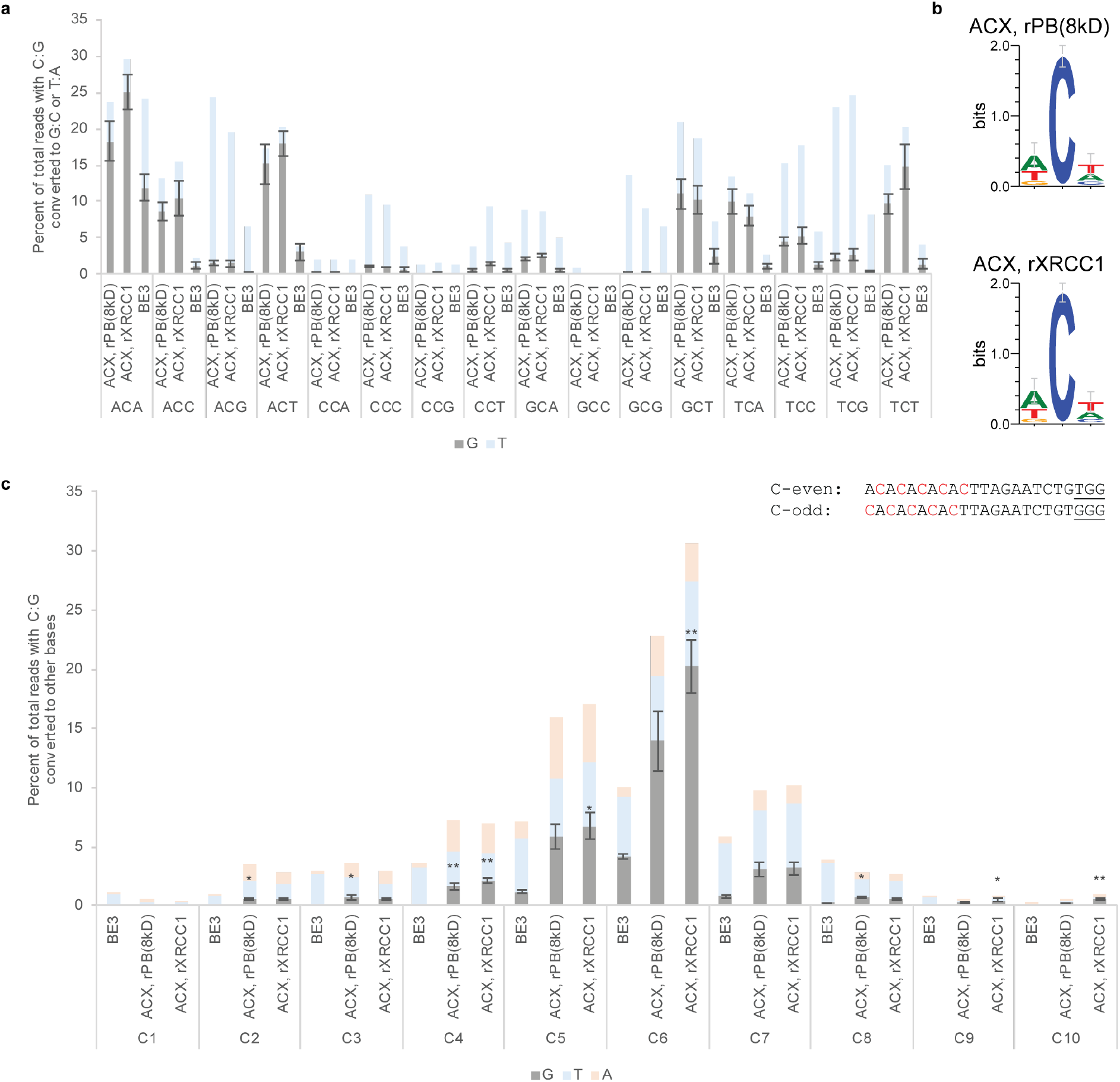
Sequence context and editing window for two selected CGBEs. **(a)** CGBEs effect C:G to G:C editing at each NCN DNA motif. gRNA sequences can be found in Extended Data Table 2. (n=2, mean ± standard error). **(b)** DNA WebLogo created with target motifs in which C:G at position 6 was edited to G:C (n=2; error bars are Bayesian 95% confidence intervals). **(c)** Editing window of CGBEs using gRNAs with alternating 5’-W-C-3’ motifs. Targeted C’s are in red. PAMs are underlined. *p < 0.05; **p < 0.01 (one-way ANOVA against ‘BE3,’ Dunn-Šidák; n=3, mean ± standard error).

We previously observed a reduction in editing levels at position 4 relative to position 6 of HEK2 (Extended Data Fig. 1). Similarly, editing levels at position 3 and position 4 of HEK3 are lower than that at position 5 (Extended Data Fig. 2). Both observations suggested that our candidates might not inherit the exact base editing window of BE3^1^. To elucidate the editing window of the CGBEs, we chose a genomic site that has an alternating 5’-W-C-W-C-W-C-3’ sequence such that gRNAs can be designed with C’s located either at every odd position or at every even position (Fig. 2c). We detected BE3 C:G to T:A editing between positions 1 and 9, with higher editing efficiency between positions 4 and 8. For both ACX, rPB(8kD) and ACX, rXRCC1, C:G to G:C editing was observed in a nine nucleotide window from positions 2 to 10. However, only at positions 5 and 6 was C:G to G:C editing the predominant and appreciable outcome.

Recognizing that simply removing UGI can potentially increase C:G to G:C editing at the expense of C:G to T:A editing^2^, we next sought to quantify the effect of fusing rXRCC1 or rPB(8kD) to rApobec-nCas9. Across 28 independent treatments using a variety of gRNAs (Fig. 3, Extended Data Fig. 5), BE3 was able to effect on average 4.4% C:G to G:C editing. Removing UGI from BE3 raised mean C:G to G:C editing to 11.5%. Fusing rPB(8kD) did not provide additional significant effect (p > 0.05). However, fusing rXRCC1 further raised mean C:G to G:C editing levels to 15.3% (p = 0.01). Meanwhile, the major byproduct C:G to T:A editing stayed between 6% and 9%. Our results suggest that UGI removal from BE3 promotes C:G to G:C editing; rXRCC1 fusion further increases C:G to G:C editing. Therefore, we narrowed down to rAPOBEC-nCas9-rXRCC1 as the preferred CGBE embodiment that effects C:G to G:C editing at 15±7% efficiency in human cells, at a 68±14% purity, within a two-nucleotide target window and WCW, ACC, and GCT target sequence contexts.

**Fig. 3.**
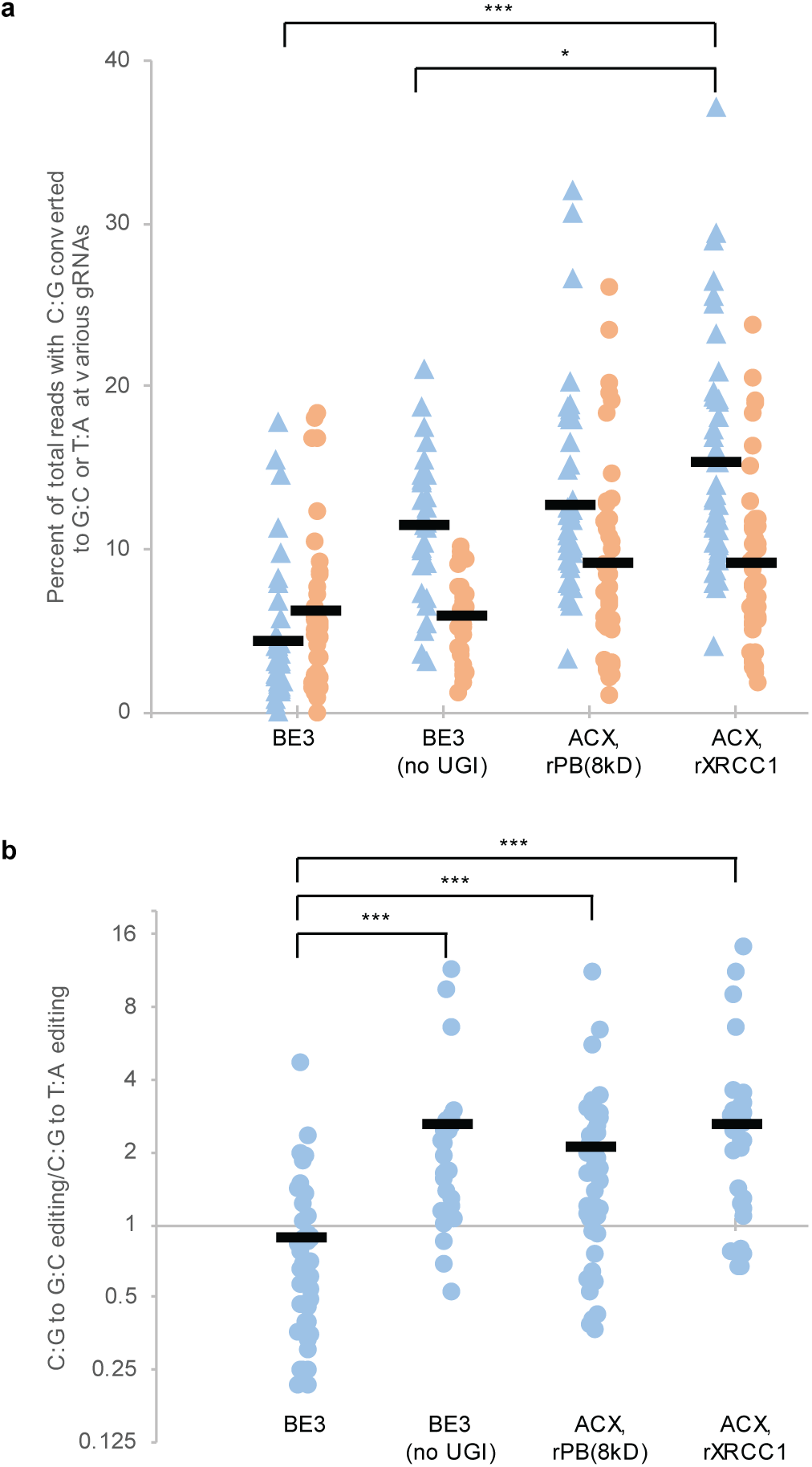
CGBE induces efficient C:G to G:C editing as the predominant product. **(a)** Removal of UGI from BE3 increases C:G to G:C editing (blue triangles); fusion of rXRCC1 further increases C:G to G:C editing. The major byproduct is C:G to T:A editing (orange circles). **(b)** Mean C:G to G:C editing/C:G to T:A editing ratio. Data includes all biological replicates across 15 genomic loci that reside within a WCW, ACC or GCT motif. *p < 0.05; ***p < 0.001 (two-tailed Student’s T test; n=28-45 biological replicates).

Here, we developed CGBEs that target cytosine within a specified window and convert it into guanine as a predominant editing product. In separate work, Liu and Koblan designed CGBE candidates by fusing UDG, UdgX, and polymerases with rAPOBEC-nCas9^12^ (Extended Data Fig. 6). Both studies induce a C to U change via rAPOBEC and envision this U to be further converted to an abasic site; but the downstream resolution of this abasic site is distinctly different. As a direct opposite of our study, which seeks to induce BER and hence repair the targeted abasic site (Extended Data Fig. 7), Liu and Koblan designed candidates to maintain the abasic site while polymerases perform translesion synthesis on the opposite strand. A similar translesion polymerization hypothesis was also put forth by Gajula^13^. This proposed mechanism is unlikely to be the case for our CGBEs. Instead, we envisage that upon creation of an abasic site in the Cas9-induced R-loop, cellular UDG is displaced by APE1, after which XRCC1 recruits various BER components^6,14^ to repair the abasic site independently of the unedited opposite strand, giving rise to guanine as the major product. Subsequent DNA repair converts the G:G mismatch to G:C. Such a hypothesis would be consistent with the tight binding of Cas9 on its target strand^15^, which renders that strand less accessible to other enzymes; the accessibility of the deaminated strand as a single-strand of the R-loop^16,17^; the detrimental effect of UDG and UdgX on C:G to G:C editing^12^ (Extended Data Fig. 6), which suggests that the persistence of an UDG-bound site or abasic site might act against the C:G to G:C reaction; and the C:G to G:C editing effected by our CGBEs with domains that do not have intrinsic polymerase activity but are key drivers of abasic site repair. While further mechanistic studies and development continue, CGBEs expand the growing suite of precise genome-editing tools that include CBEs^1,2^, ABEs^3^, and prime editors^16^, together enabling the precise and efficient engineering of DNA for biological interrogation and disease correction.

## Methods

### Constructs and cell culture

DNA amplification was done via PCR using Q5 Hot Start HiFi 2X Master Mix (NEB, M0494). All plasmids were constructed using either blunt end ligation or Gibson Assembly. For blunt end ligation, PCR products were first treated with DpnI (NEB, R0176) and T4 Polynucleotide Kinase (NEB, M0201) before being ligated to plasmid vectors using T4 DNA Ligase (NEB, M0202). NEBuilder HiFi DNA Assembly Master Mix (NEB, E2621) was used for Gibson Assemblies. hXRCC1 (pTXG-hXRCC1) and hLIG3 (pGEX4T-hLIG3) were gifts from Primo Schaer (Addgene plasmid # 52283 and # 81055 respectively). BE3 (pCMV-BE3) was a gift from David R. Liu (Addgene plasmid # 73021). The mutation R400Q is introduced to hXRCC1; N628K is introduced to hLIG3. rXRCC1, rLIG3, hPBs, and rPBs were obtained as human codon-optimized de novo synthesized gene fragments (Twist Biosciences). All other oligonucleotides used in the study were *de novo* synthesized (IDTDNA).

Following transformation of constructs and bacterial outgrowth, gRNA plasmids and base-editor plasmids were purified using the PureYield Plasmid Miniprep System (Promega, A1223) and Plasmid Plus Maxi Kit (Qiagen, 12965) respectively. Both plasmid kits’ protocols include endotoxin removal steps. Plasmids were Sanger sequenced for sequence verification.

### Cell culture, transfection, and genomic DNA harvest

HEK293AAV cells (Agilent, 240073) were maintained in DMEM (Thermo Fisher, 10569-010) supplemented with 10% HI FBS (Thermo Fisher). 30,000 HEK293AAV cells were added to each well of a 48 well plate one day before transfection. For each well, 750 ng of base-editor plasmid, 250 ng of gRNA plasmid, and 20 ng of GFP plasmid were transfected into these cells using Lipofectamine 3000 (Invitrogen, L3000015) according to the manufacturer’s protocol. The media were replaced with fresh media one day after transfection. Three days after transfection, media were removed, cells were washed with 50 μL PBS, pH 7.2 (Thermo Fisher, 20012-027), and genomic DNA was extracted using 50 μL of Quick Extract DNA Extract Solution (Lucigen, QE09050) per well according to manufacturer’s protocol. All sample sizes indicate biological replicates.

### Sequencing of genomic DNA

Sites of interest were prepared for high-throughput sequencing via two PCR amplifications – the first PCR amplifies the region of interest while the second PCR adds appropriate sequencing barcodes. Primers for the first PCR are included in Extended Data Table 3. Primers for the second PCR are based off Illumina adaptors. Amplicons from the second PCR were then pooled and gel extracted (Promega, A9282) to make the final library, which was quantified via Qubit fluorometer (Thermo Fisher) and sequenced on an Illumina iSeq 100 according to the manufacturer’s protocol. The resultant FASTQ files were analyzed using CRISPResso2^18^. All sample sizes indicate biological replicates.

Statistical analyses were performed on Matlab. Weblogos were created using Weblogo 3^19^.

## Data availability

High-throughput sequencing data will be deposited to the NCBI Sequence Read Archive database. Plasmid encoding ACX, rXRCC1 will be available on Addgene.

## Acknowledgements

We thank Anna Traczyk for helpful discussions; Eddie Keng for help with HTS; Hannah Nicholas, Nicholas Ong, and Hong-Ting Prekop for editing; and Ke Guo and Daryl Lim for general administrative support. This work is supported by Agency for Science, Technology and Research (A*STAR) and Industrial Alignment Fund Pre-Positioning (IAF-PP) grant H17/01/a0/012. Work by P.D.R. and W.L.C. is further supported by Advanced Manufacturing and Engineering (AME) Programmatic grant A18A9b0060.

## Author contributions

L.C. made the initial discovery. L.C., J.E.P., and W.L.C. designed the research. L.C., J.E.P., and P.P. performed the experiments. P.D.R. performed biological replicates. Y.T.C. and S.N.M. cloned plasmids. L.C., J.E.P., P.P., and W.L.C. analyzed data. L.C. and W.L.C. wrote the manuscript with input from other authors. W.L.C. supervised the research.

## Additional information

Supplementary Information is available for this paper.

Correspondence and requests for materials should be addressed to W. L. C.

## Extended data figures, tables and legends

**Extended Data Fig. 1.**
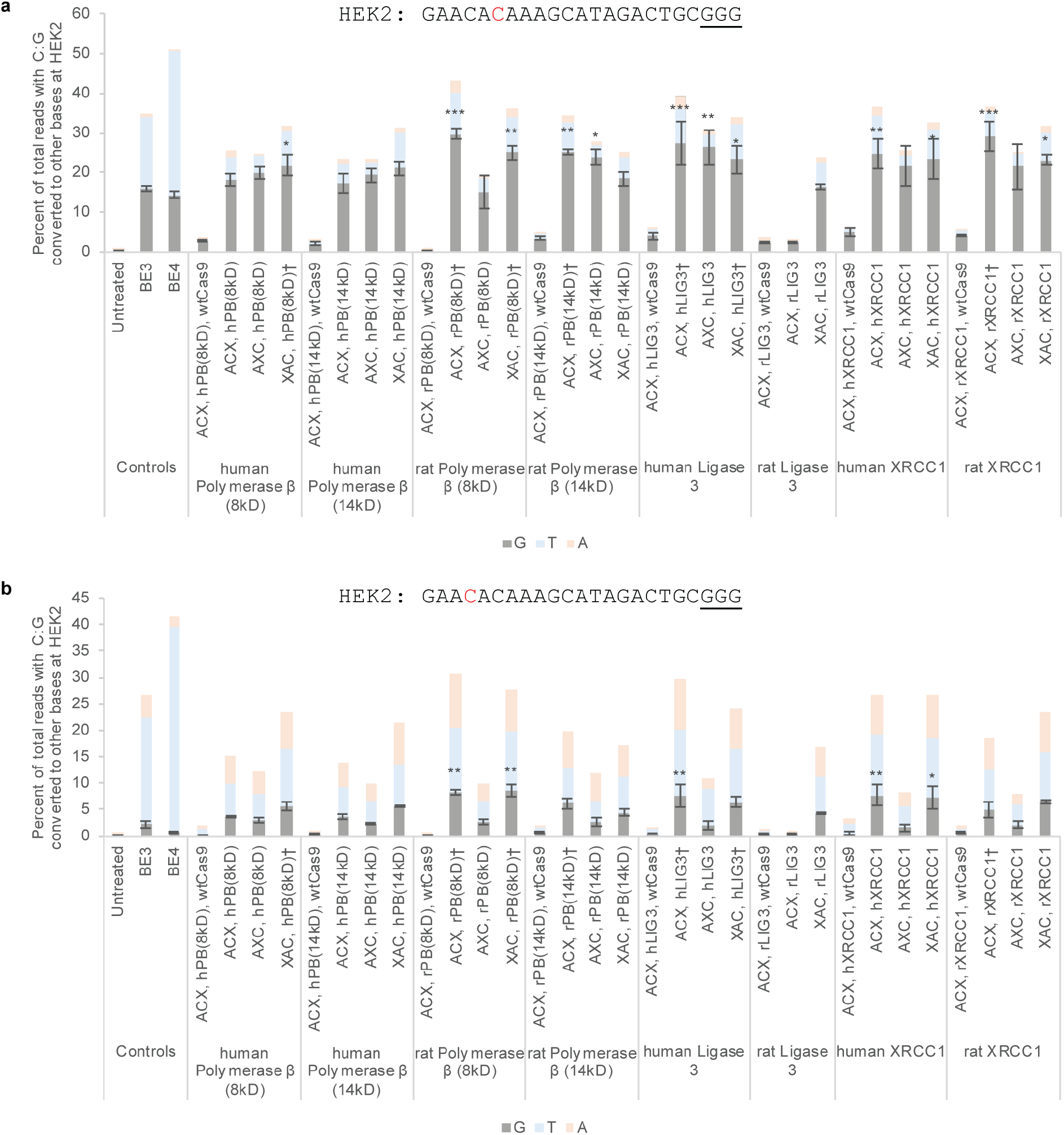
Initial screen of CGBE candidates on HEK2. **(a)** For some CGBE candidates, C:G to G:C editing is the predominant edit at position 6. **(b)** C:G to T:A editing is the predominant edit at position 4. The seven candidates selected for further studies are marked with †. Targeted C’s are in red. PAMs are underlined. *p < 0.05; **p < 0.01; ***p < 0.001 (one-way ANOVA against ‘Untreated,’ Dunn-Sidak; n=3, mean ± standard error).

**Extended Data Fig. 2.**
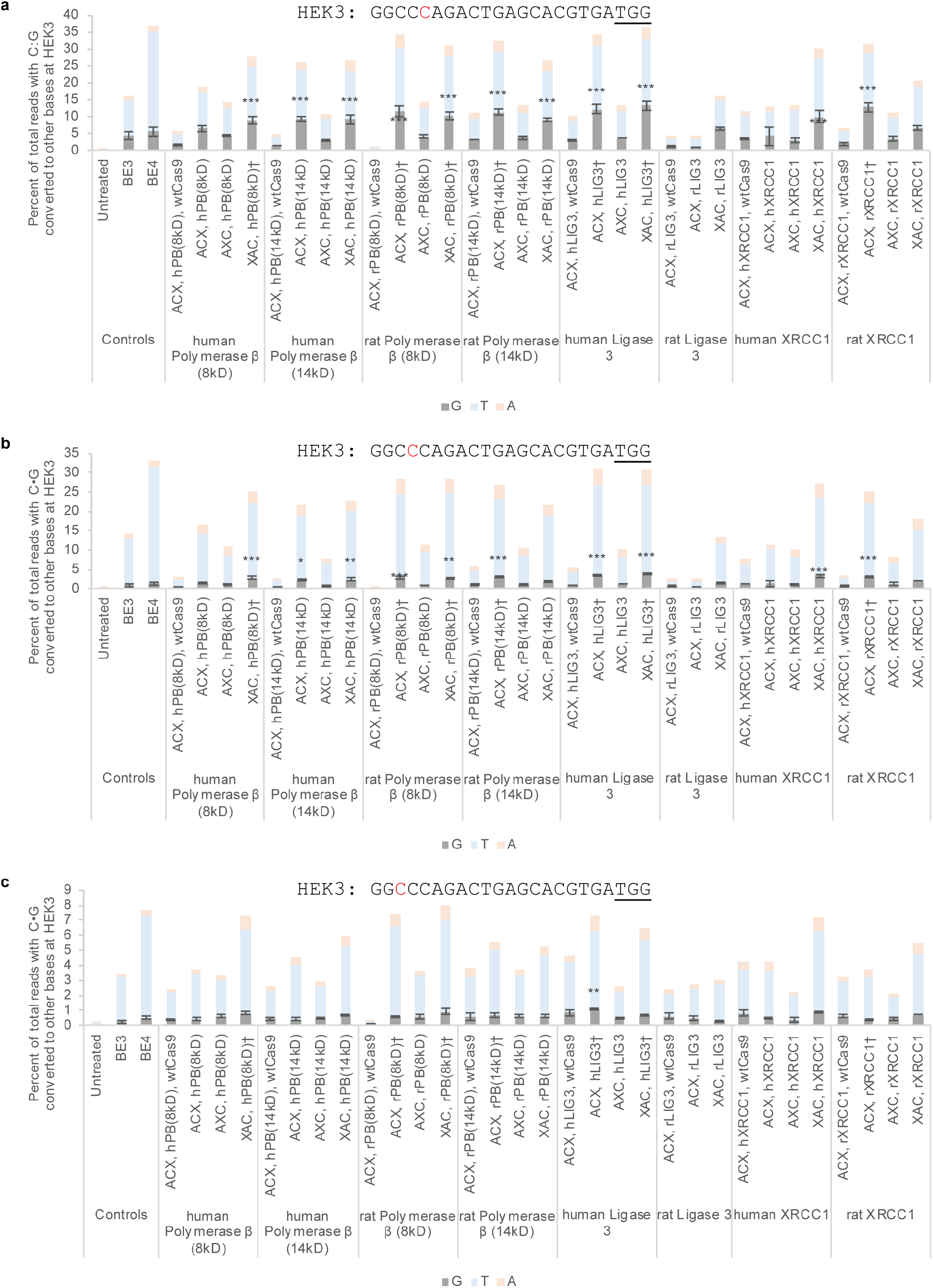
Initial screen of CGBE candidates on HEK3 at (a) position 5, (b) position 4, and (c) position 3. The seven candidates selected for further studies are marked with †. Targeted C is in red. PAM is underlined. *p < 0.05; **p < 0.01; ***p < 0.001 (one-way ANOVA against ‘Untreated,’ Dunn-Šidák; n=3, mean ± standard error).

**Extended Data Fig. 3.**
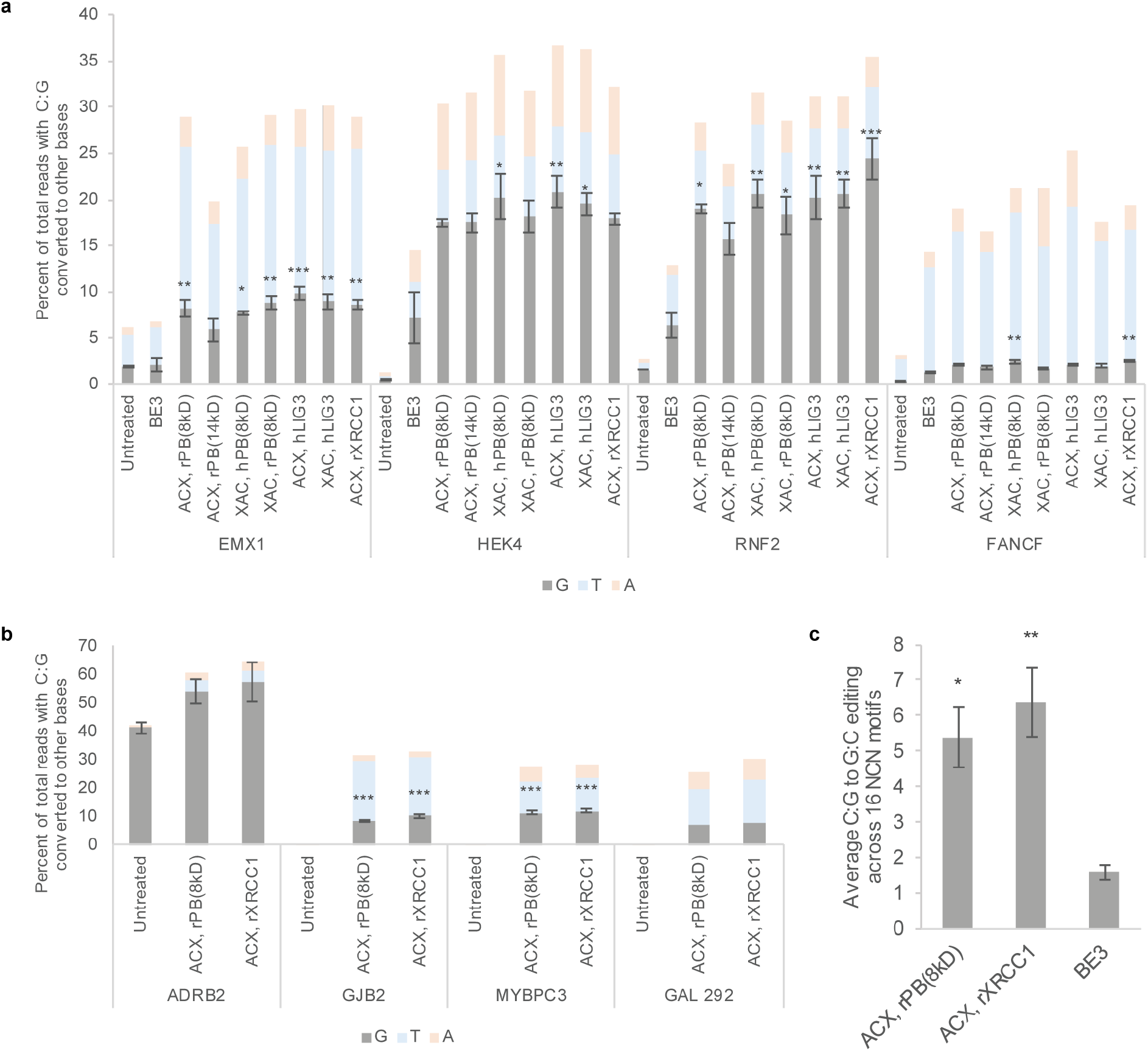
Further characterization of shortlisted CGBE candidates. **(a)** CGBE candidates effect C:G to G:C mutations at EMX1, HEK4, RNF2, and FANCF. C:G to G:C editing is the main edit at HEK4 and RNF2; C:G to T:A editing is the main edit at FANCF and EMX1. (one-way ANOVA against ‘BE3,’ Dunn-Šidák; n=3, mean ± standard error) **(b)** CGBE editing at disease-associated genes ADRB2, GJB2, MYBPC3, and GAL 292. Note that ADRB2 contains naturally occurring polymorphism in HEK293AAV cells, and hence this data is not included in Fig 3. (one-way ANOVA against ‘Untreated,’ Dunn-Šidák; n=5 for ADRB2 and MYBPC3; n=4 for GJB2; n=1 for GAL 292, mean ± standard error) **(c)** Mean C:G to G:C editing as a percent of all reads across 16 NCN sites. CGBEs increase C:G to G:C editing by three to four fold compared to BE3, across all possible NCN sequences. gRNA sequences are included in Extended Data Table 2. *p < 0.05; **p < 0.01; ***p < 0.001 (one-way ANOVA against ‘BE3,’ Dunn-Šidák; n=32, mean ± standard error).

**Extended Data Fig. 4.**
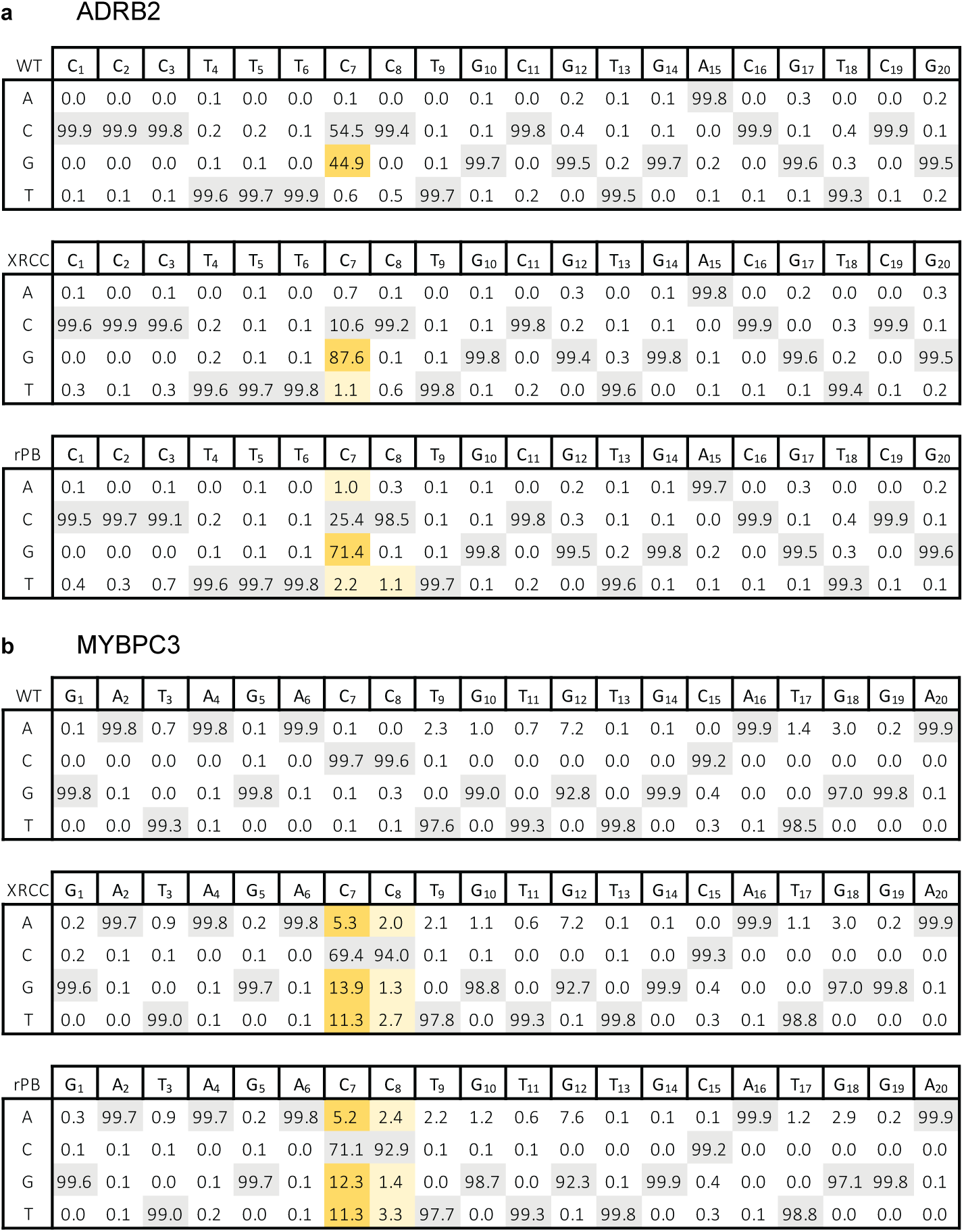
Representative data of CGBE editing at (a) ADRB2 and (b) MYBPC3. WT denotes wild-type untreated cells; XRCC denotes ACX, XRCC1; rPB denotes ACX, rPB(8kD). Note that ADRB2 contains naturally occurring polymorphism in HEK293AAV cells.

**Extended Data Fig. 5.**
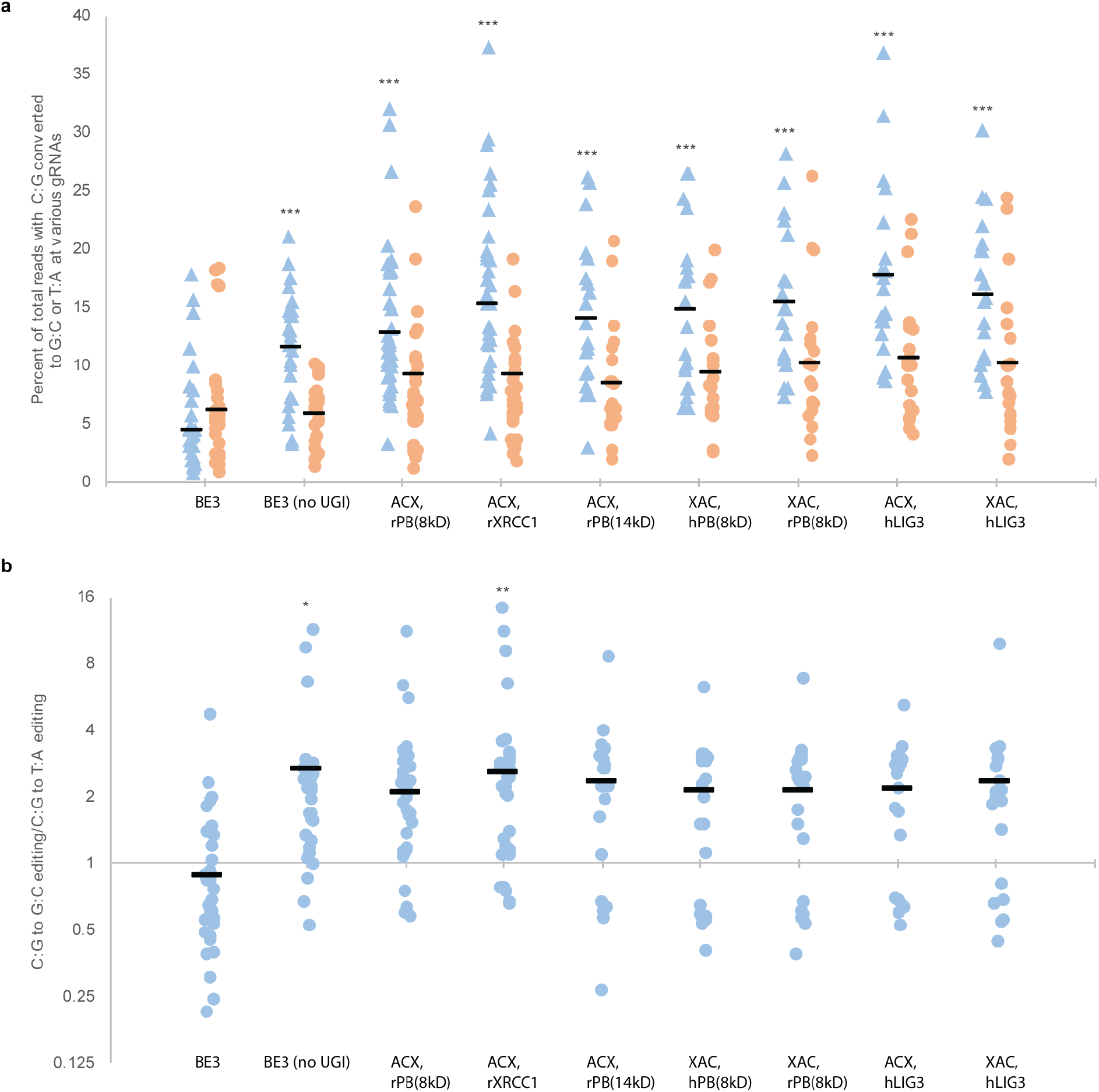
ACX, rXRCC1 is the best performer out of shortlisted CGBE candidates. **(a)** C:G to G:C editing (blue triangles) vs. C:G to T:A editing (orange circles) as percent of all reads across gRNAs used in this study. All biological replicates are included except those targeting the 10 suboptimal C:G to G:C base editing motifs (Fig. 2a and Fig. 2b) and ADRB2 due to naturally occurring polymorphism (Extended Data Fig. 3b). **(b)** Ratio of C:G to G:C editing to C:G to T:A editing across gRNAs used in this study. Only BE3 (no UGI) and ACX, rXRCC1 give a significantly higher ratio of C:G to G:C editing/C:G to T:A editing. *p < 0.05; **p < 0.01; ***p < 0.001 (one-way ANOVA against ‘BE3,’ Dunn-Šidák, n=20-45 biological replicates).

**Extended Data Fig. 6.**
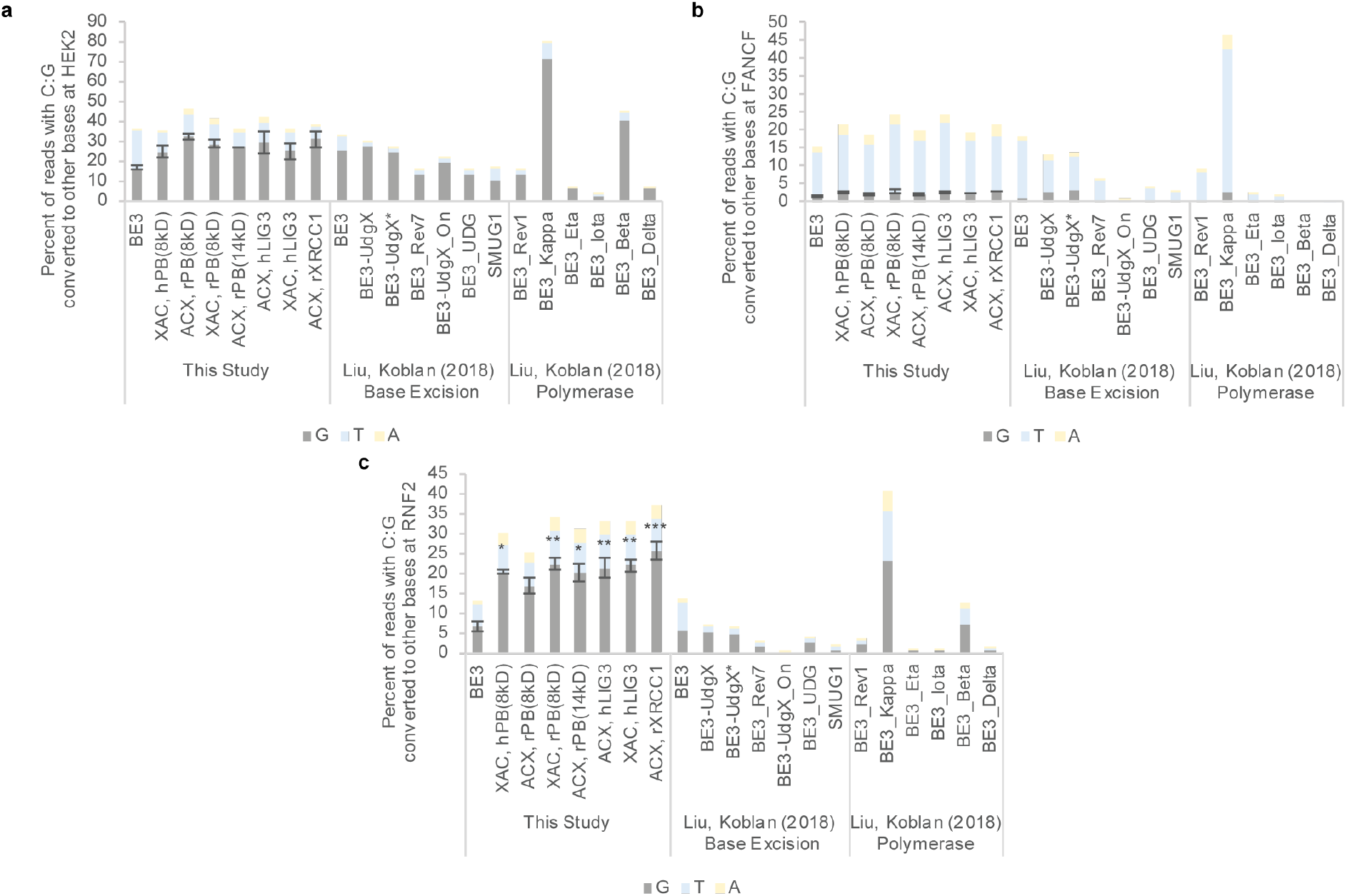
Comparison of CGBEs at (a) HEK2, (b) FANCF, and (c) RNF2. Since datasets were generated independently and in different cell types, comparisons should be made only against BE3 common to the two studies. Fusion of base excision enzymes such as UDG and UdgX decreases C:G to G:C editing beyond BE3 at 2 of 3 sites (n=1; Liu and Koblan, 2018). Fusion of base excision repair enzymes such as rXRCC1 increases C:G to G:C editing beyond BE3. *p < 0.05; **p < 0.01; ***p < 0.001 (one-way ANOVA against ‘BE3,’ Dunn-Šidák; n=3, mean ± standard error; this study).

**Extended Data Fig. 7.**
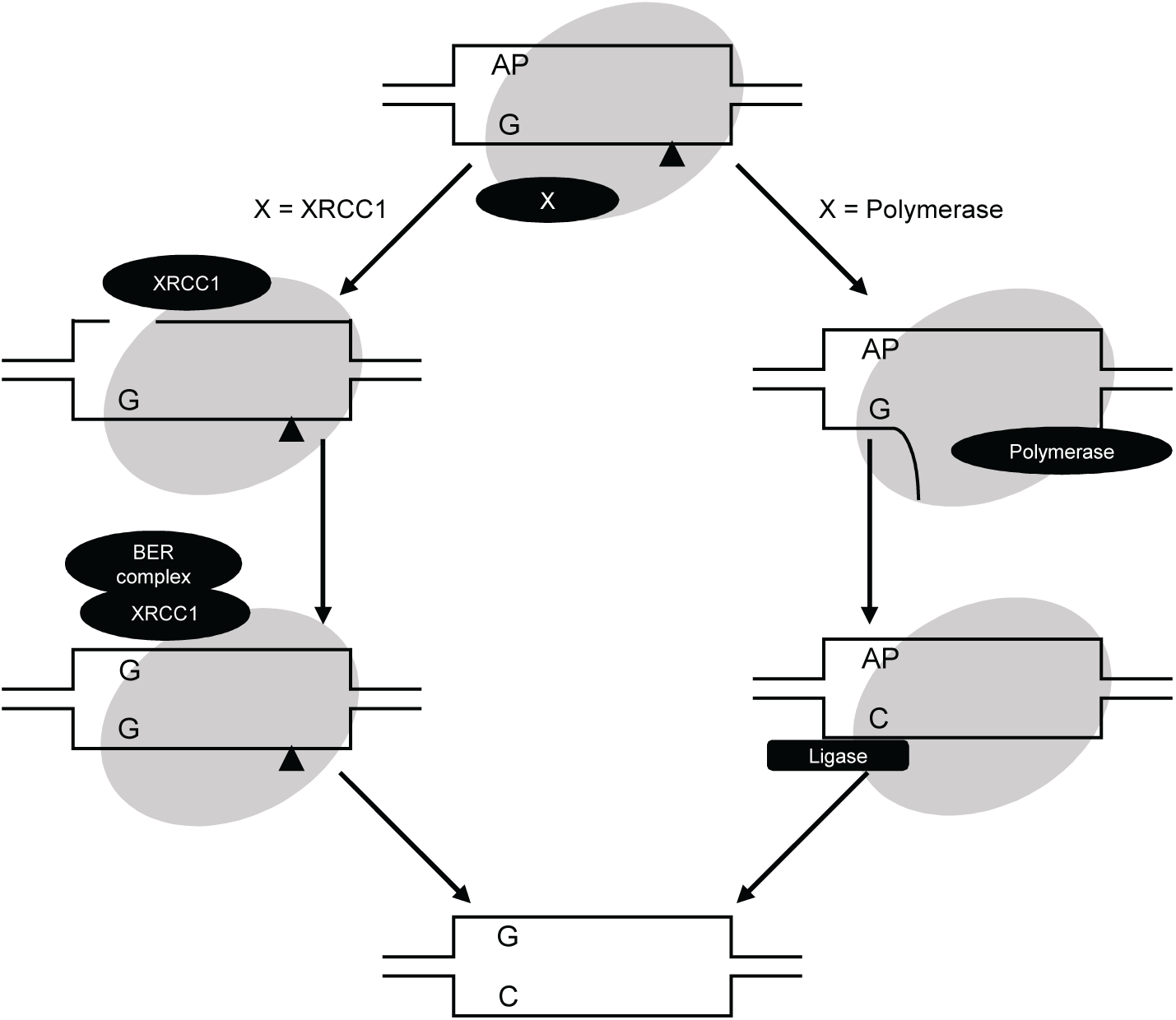
Distinct strategies for CGBE design. This study employs a CGBE design strategy (left) where Cas9 is fused to protein(s) involved in repairing uracil-containing or abasic sites (AP). The activities of these BER proteins are expected to convert the AP to G before the nucleotide on the opposite strand is converted from G to C. In contrast, the strategy employed by Liu and Koblan (right) seeks to maintain the abasic site (AP) throughout a translesion synthesis envisioned to occur on the opposite strand. The AP is repaired after the nucleotide on the opposite strand is converted from G to C. In other words, this study employs proteins that repair and not maintain abasic sites whereas Liu and Koblan employ proteins that maintain and not repair abasic sites.

**Extended Data Table 1.** CGBEs enable treatment avenues to previously unaddressable SNPs that represent half of all known human disease-associated SNPs. CBE enables treatment to 48% of all known disease-associated SNPs, while ABE enables treatment to 6%. CGBEs effect primarily C:G to G:C and G:C to C:G changes (*yellow highlight*) that can correct 11% of disease-associated SNPs. In combination with CBEs or ABEs, CGBEs effect secondarily G to T, C to A, A to C, and T to G edits (*green highlight*). With CBEs, ABEs, and CGBEs, the remaining 7% of SNPs (A to T and T to A) can also be corrected (*orange highlight*).

| From | To | % of SNPs in ClinVar | Equivalent to |  | Base-editing route for correction |
| --- | --- | --- | --- | --- | --- |
| A | T | 7 | T | A | A to G (ABE), G to C (CGBE), C to T (CBE) |
|  | G | 48 |  | C | ABE |
|  | C | 15 |  | G | C to G (CGBE), G to A (CBE) |
| G | A | 6 | C | T | CBE |
|  | T | 14 |  | A | A to G (ABE), G to C (CGBE) |
|  | C | 11 |  | G | CGBE |

**Extended Data Table 2.**
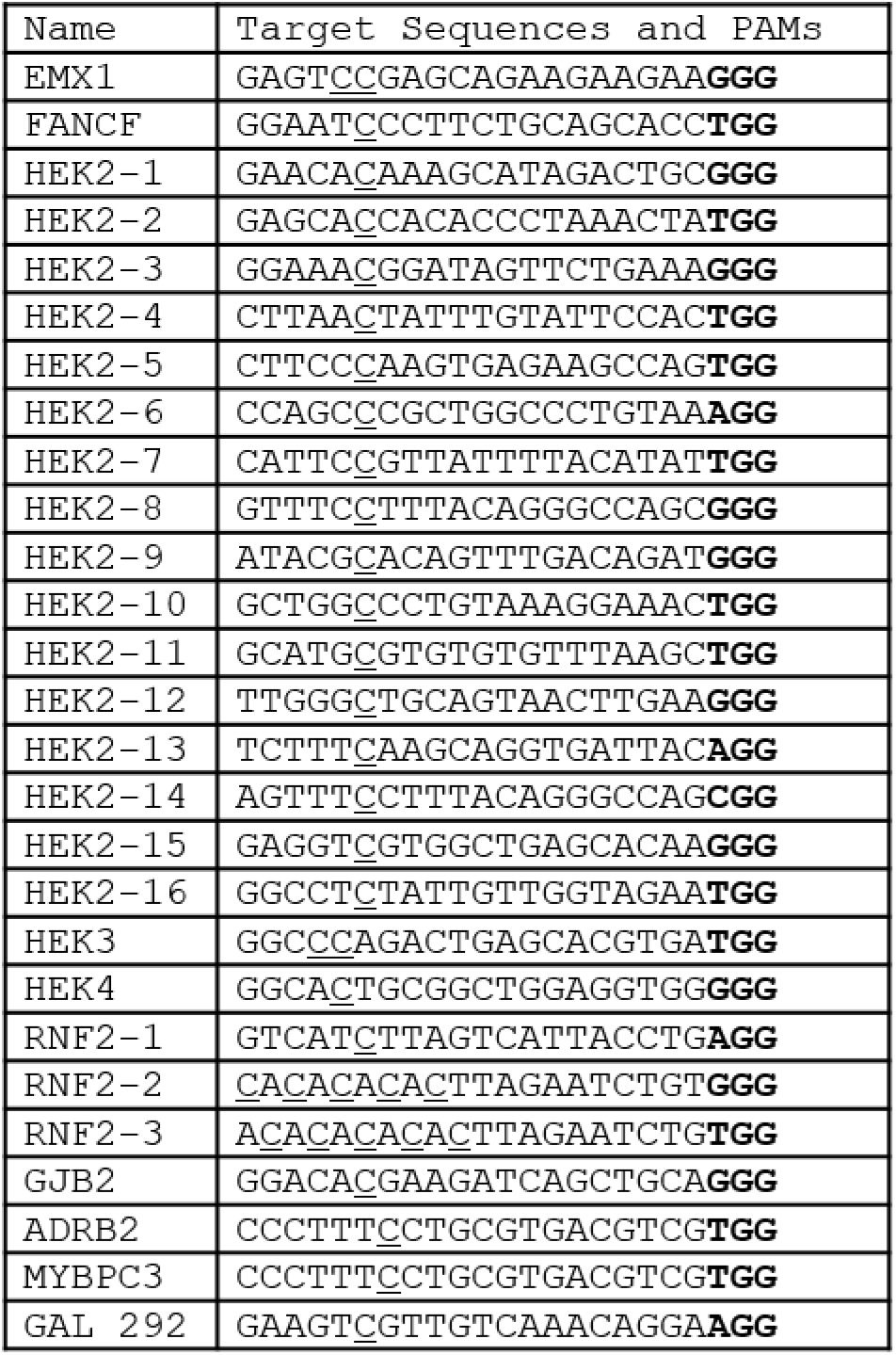
Target protospacer sequences used in this study. Targeted C’s are underlined. PAMs are in bold.

